# Two Lytic Bacteriophages That Depend on the *Escherichia coli* Multi-Drug Efflux Gene *tolC* and Differentially Affect Bacterial Growth and Selection

**DOI:** 10.1101/397695

**Authors:** Alita R. Burmeister, Rose G. Bender, Abigail Fortier, Adam J. Lessing, Benjamin K. Chan, Paul E. Turner

## Abstract

Bacterial pathogens are increasingly evolving drug resistance under natural selection from antibiotics in medicine, agriculture, and nature. Meanwhile, bacteria ubiquitously encounter bacteriophages and can rapidly evolve phage resistance. However, the role of phages in interacting with drug-resistant and drug-sensitive bacteria remains unclear. To gain insight into such relationships, we screened for and characterized phages that rely on the multi-drug efflux pump gene *tolC*. First, we screened a collection of 33 environmental and commercial *Escherichia coli* phages for their ability to infect cells that lacked *tolC*. Our screen revealed two phages that had reduced efficiency of plating (EOP) on the *tolC* knockout compared to wild type. We further characterized these phages with bacterial growth curves, transmission electron microscopy, and analysis of phage-resistant mutants. Phage U136B is a curly-tailed virus in family *Siphoviridae* with no ability to infect a *tolC* knockout, suggesting TolC is the U136B receptor. Phage 132 is a contractile-tailed virus in family *Myoviridae* with reduced EOP on cells lacking *ompF* and its positive regulators *tolC* and *ompR*. U136B and 132 differentially effect bacterial growth and lysis, and U136B-resistant mutants contain mutations of the *tolC* gene. Together, these results show that the *tolC* gene involved in drug resistance can modify bacteria-phage interactions in multiple ways, altering bacterial lysis and selection. These new phages offer utility for studying evolution, tradeoffs, and infection mechanisms.

**Importance:** Bacteria face strong selection by antibiotics in medicine and agriculture, resulting in increasing levels of drug resistance among bacterial pathogens. Slowing this process will require an understanding of the environmental contexts in which drug resistance evolutionarily increases or decreases. In this study, we investigate two newly-isolated bacteriophages that rely on a bacterial antibiotic resistance gene. These bacteriophages vary in their interactions with drug-resistant bacteria, with one of the phages selecting for phage-resistant mutants that have mutations in the antibiotic resistance gene. Further study of these new phages will be useful to understanding evolutionary tradeoffs and how phages might be applied in natural settings to reverse the problem of drug resistance.

## Introduction

Widespread use of antibiotics in medicine and agriculture has selected for the evolution of multi-drug resistant (MDR) bacterial pathogens (1). Meanwhile, bacteria frequently encounter phages, which are prevalent in the human microbiota, in hospital and farm settings, and in natural environments (2), and which exert selection pressure for bacteria to resist phage exploitation (3-7). However, the interaction between selection from antibiotics and phages, along with its role in driving bacterial evolution, remain unclear, in part because these interactions depend on both the environment and specific phage species.

Potential evolutionary interactions between drug resistance and phage resistance mechanisms in bacteria have been previously identified, including both positive and negative interactions that are highly genotype-dependent (8, 9). For example, *Pseudomonas aeruginosa* bacteria that evolve resistance to phage 14/1 simultaneously become *more* resistant to antibiotics (10), whereas *P. aeruginosa* that evolve resistance to phage OMKO1 become *less* resistant to antibiotics (5). In *Escherichia coli*, bacteria that evolve resistance to phage TLS also lose antibiotic resistance (11). Such interactions demonstrate that multiple selection pressures sometimes cause bacteria to evolve mutations with *trade-up* potential, whereby phages contribute to the problems of increased antibiotic resistance and virulence; in other cases, the mutations have *trade-off* potential, whereby phages reduce the problem of antibiotic resistance.

Bacteria-phage interactions can be highly dependent on cell membrane proteins. Such proteins are often exploited by phages for cell attachment and entry. In particular, multi-drug efflux pumps are protein complexes spanning the inner and outer membranes of some bacteria, such as the homologous TolC-AcrAB system in *Escherichia coli* and OprM-MexAB system in *P. aeruginosa* (12). These efflux systems confer resistance to multiple antibiotics, acting as generalized transporters for multiple antibiotic classes as well as detergents, dyes, and bile acids (13). The outer membrane protein (OMP) components (TolC or OprM) are membrane-spanning beta barrels, with peptide loops that extend outside of the cell. The extracellular loops of OMPs are frequently exploited by phages as the specific binding sites for initiating phage infection (11, 14-16). When phages use these OMPs as receptors, bacteria face selection for reduced or modified OMPs, catalyzing ecological restructuring or coevolutionary arms races that in turn alter selection on the phages (6, 15, 17, 18). Additionally, loss or modification of OMP genes has been shown to alter expression of other OMP genes. For example, *tolC* mutants have reduced expression of outer membrane proteins OmpF, NmpC, and protein 2 (19). Therefore, loss of an OMP gene might impact a phage either directly – by loss of the phage receptor – or indirectly, through changes to the expression of the phage receptor.

In this study, we sought out phages that rely on the antibiotic resistance gene *tolC*, which encodes the outer membrane protein of the TolC-AcrAB efflux pump. Such phages, like the previously-characterized TolC-targeting phage TLS (11), might impose selection on bacterial communities to evolve phage resistance while losing antibiotic resistance. Such phages will be useful to the laboratory study of evolutionary tradeoffs, or more practically, to restore drug sensitivity in clinical settings. To search for such phages, we conducted a screen of our *E. coli* phage library on bacteria that lacked the *tolC* gene. Out of 33 phages, we found two with reduced plaquing efficiency on the *tolC* knockout. We found that these phages differentially affect both bacterial population dynamics and the potential for evolution of antibiotic resistance.

## Results

### Phage interactions with *tolC*^−^ bacteria

We conducted a screen of newly-collected phages with unknown receptors by plating for plaques on both wild-type *E. coli* and its isogenic *tolC* knockout from the Keio collection (20) (Table 1). Of 33 phages, we found two (phage U136B and phage 132) with dramatically reduced efficiency of plating (EOP, the number of plaques formed on a mutant strain of bacteria relative to a wild type) (Fig. S1, Table 1). Phage U136B, isolated from a swine farm in Connecticut, appears to obligately require bacterial *tolC* for infection, with EOP below our detection limit of 10^-10^ (data not shown); we have never observed a plaque of U136B in the absence of the *tolC* gene. Phage 132, also isolated from a swine farm in Connecticut, has dramatically reduced plating efficiency on *tolC*^−^ (EOP = 9.5×10^-6^) when plated with the standard Luria Bertani (LB) top agar formula (7.5 g/L agar). The phage 132 EOP on *tolC*^−^ bacteria increases – but does not fully recover – when the top agar contains only 3.8 g/L agar, which we discuss in the Results section.

**Table 1.** *E. coli* and bacteriophage strains used in this study. All bacteria were obtained from the Yale Coli Genetic Stock Center and were originally described by Baba *et al.* 2008 (20).

| Strain | Description | Relevant Characteristics |
| --- | --- | --- |
| **Bacteria** |  |  |
| BW25113 | *E. coli* K12, parental strain of Keio collection |  |
| JW5503-1 | *tolC* knockout in Keio collection | $\Delta tolC732::kan$ |
| JW0451-2 | *acrB* knockout in Keio collection | $\Delta acrB747::kan$ |
| JW0452-3 | *acrA* knockout in Keio collection | $\Delta acrA748::kan$ |
| JW2341-1 | *fadL* knockout in Keio collection | $\Delta fadL752::kan$ |
| JW0401-1 | *tsx* knockout in Keio collection | $\Delta tsx773::kan$ |
| JW3996-1 | *lamB* knockout in Keio collection | $\Delta lamB732::kan$ |
| JW0940-6 | *ompA* knockout in Keio collection | $\Delta ompA772::kan$ |
| JW2203-1 | *ompC* knockout in Keio collection | $\Delta ompC768::kan$ |
| JW0146-2 | *fhuA* knockout in Keio collection | $\Delta fhuA766::kan$ |
| JW5195-1 | *tonB* knockout in Keio collection | $\Delta tonB760::kan$ |
| JW0912-1 | *ompF* knockout in Keio collection | $\Delta ompF746::kan$ |
| JW3368-1 | *ompR* knockout in Keio collection | $\Delta ompR739::kan$ |
| JW3603-2 | *rfaB* knockout in Keio collection | $\Delta rfaB739::kan$ |
| JW3596-1 | *rfaC* knockout in Keio collection | $\Delta rfaC733::kan$ |
| JW3594-1 | *rfaD* knockout in Keio collection | $\Delta rfaD731::kan$ |
| JW3024-1 | *rfaE* knockout in Keio collection | $\Delta rfaE745::kan$ |
| JW3595-2 | *rfaF* knockout in Keio collection | $\Delta rfaF732::kan$ |
| JW3606-1 | *rfaG* knockout in Keio collection | $\Delta rfaG742::kan$ |
| JW3818-1 | *rfaH* knockout in Keio collection | $\Delta rfaH783::kan$ |
| JW3602-1 | *rfaI* knockout in Keio collection | $\Delta rfaI738::kan$ |
| JW3601-3 | *rfaJ* knockout in Keio collection | $\Delta rfaJ737::kan$ |
| JW3597-1 | *rfaL* knockout in Keio collection | $\Delta rfaL734::kan$ |
| JW3605-1 | *rfaP* knockout in Keio collection | $\Delta rfaP741::kan$ |
| JW3607-2 | *rfaQ* knockout in Keio collection | $\Delta rfaQ743::kan$ |
| JW3604-2 | *rfaS* knockout in Keio collection | $\Delta rfaS740::kan$ |
| JW3600-1 | *rfaY* knockout in Keio collection | $\Delta rfaY736::kan$ |
| JW3599-1 | *rfaZ* knockout in Keio collection | $\Delta rfaZ735::kan$ |
| **Phage** |  |  |
| Phage 132 | Environmental isolate from swine farm |  |
| Phage U136B | Environmental isolate from swine farm |  |
| Phage T2 | Obtained from the Yale Coli Genetic Stock Center, CGSC #12141 |  |

### Phage interactions with other drug efflux pump genes

Finding that U136B and 132 rely fully or partially on presence of the *tolC* gene, we speculated that they might also require *acrA* and *acrB*, which encode the other components of the TolC multi-drug efflux pump. We performed spot tests for plaquing on the *acrA* and *acrB* knockouts of the Keio collection and found no change in plaquing ability (data not shown).

### Phage morphology

We used transmission electron microscopy to determine the general structure and morphological families of U136B and 132. Phage U136B has a curly, non-contractile tail of the *Siphoviridae* family type. It has a capsid width of 59 nm, capsid length of 61 nm, and tail length of 115 nm (Fig. 1A). Phage 132 has a contractile tail of the *Myoviridae* family type. It has a capsid width of 72 nm, capsid length of 99 nm, extended tail length of 111 nm and contracted tail length of 58 nm (Fig. 1B). Consistent with non-enveloped particle morphology, we also found that both phages were insensitive to chloroform treatment (Fig. S2). While both phages did have tails, they also appeared to be structurally robust with minimal loss due to mechanical agitation *via* vortexing (Fig. S2).

**Figure 1.**
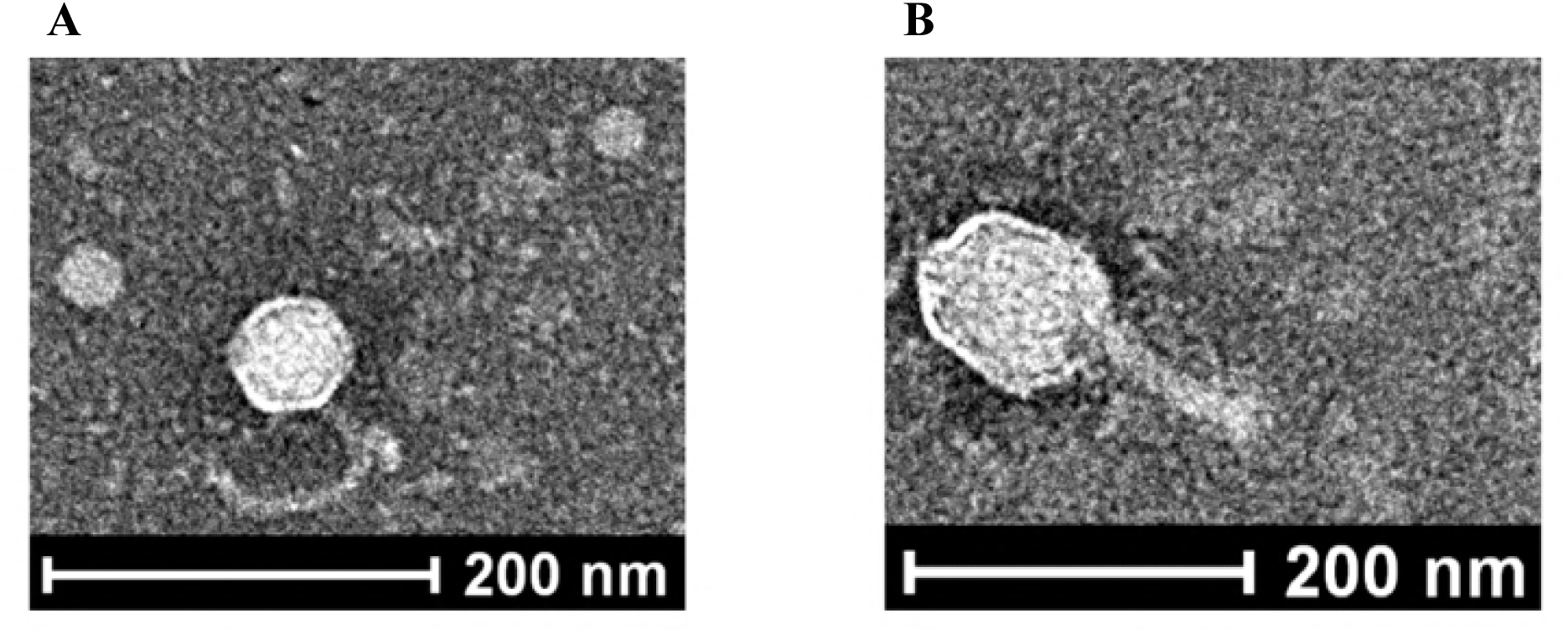
Transmission electron microscopy of the newly-isolated, *tolC*-dependent phages. A) Phage U136B. B) Phage 132.

### Phage impact on bacterial growth

The *tolC*^−^ efficiency of plating data suggested that these phages should also differentially affect the survival of wild-type and *tolC*^−^ bacteria in liquid cultures. To test this, we generated growth curves of wild-type and *tolC* knockout bacteria with and without addition of each phage. Indeed, the wild-type bacteria were killed by both phages (Fig. 2, black dashed lines), while the *tolC* knockout was unaffected by phage U136B (Fig. 2A, gray lines). Phage 132, which plaques with reduced efficiency on the *tolC* knockout, was able to kill both the *tolC* knockout and the wild-type bacteria (Fig. 2B, dashed lines).

**Figure 2.**
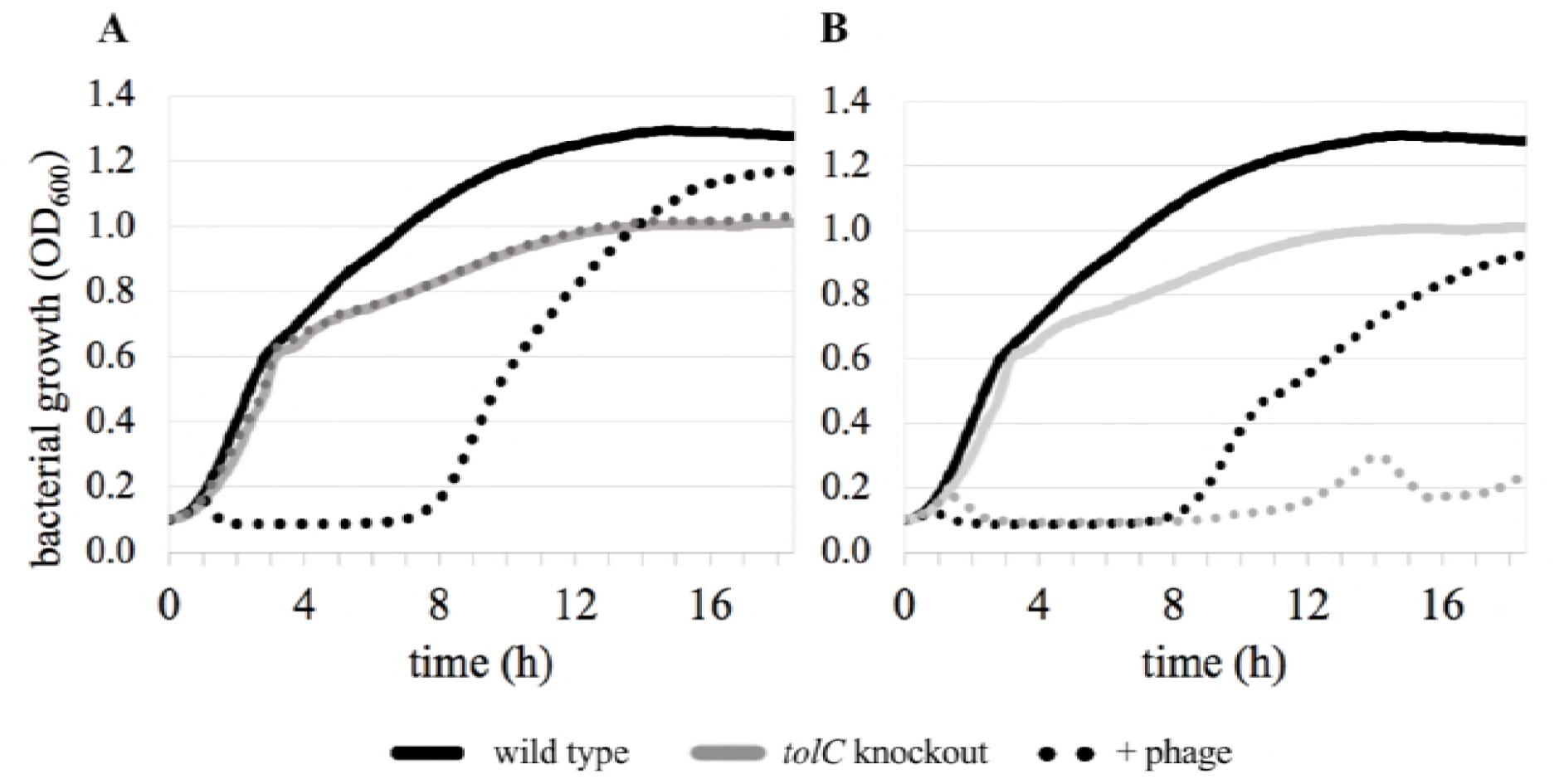
Effects of *tolC*-dependent phages U136B and 132 on bacterial growth in liquid culture. A) Knocking out *tolC* protects *E. coli* from infection by phage U136B, while the wild-type is rapidly lysed. B) Both the *tolC* knockout and wild-type are lysed by phage 132. Each curve shows the mean of three to six replicates. Individual replicates are shown in Fig. S3.

### Effect of multiplicity of infection on bacterial lysis

The ability for both phage U136B and phage 132 to rapidly lyse wild-type *E. coli* increases with higher multiplicity of infection (MOI, the ratio of phage particles to bacterial cells). Cell lysis still is rapid and efficient for both phages, beginning within two hours even at a low MOI of 0.01 (Fig. 3A-B). Increasing MOI significantly decreases the bacterial density at the onset of observed lysis (Fig. 3C), decreases the time to onset of lysis (Fig. 3D), and decreases the time to complete lysis (Fig. 3E) (p < 0.01 in all cases, Table S2 ‘MOI’). The phage type significantly affects the bacterial density at the onset of lysis and the time to complete lysis (p < 0.01 in both cases, Table S2 ‘Phage’) but not the time of lysis onset (p = 0.15). At lower MOIs, phage 132 has longer times to complete lysis than phage U136B (Fig. 3E).

**Figure 3.**
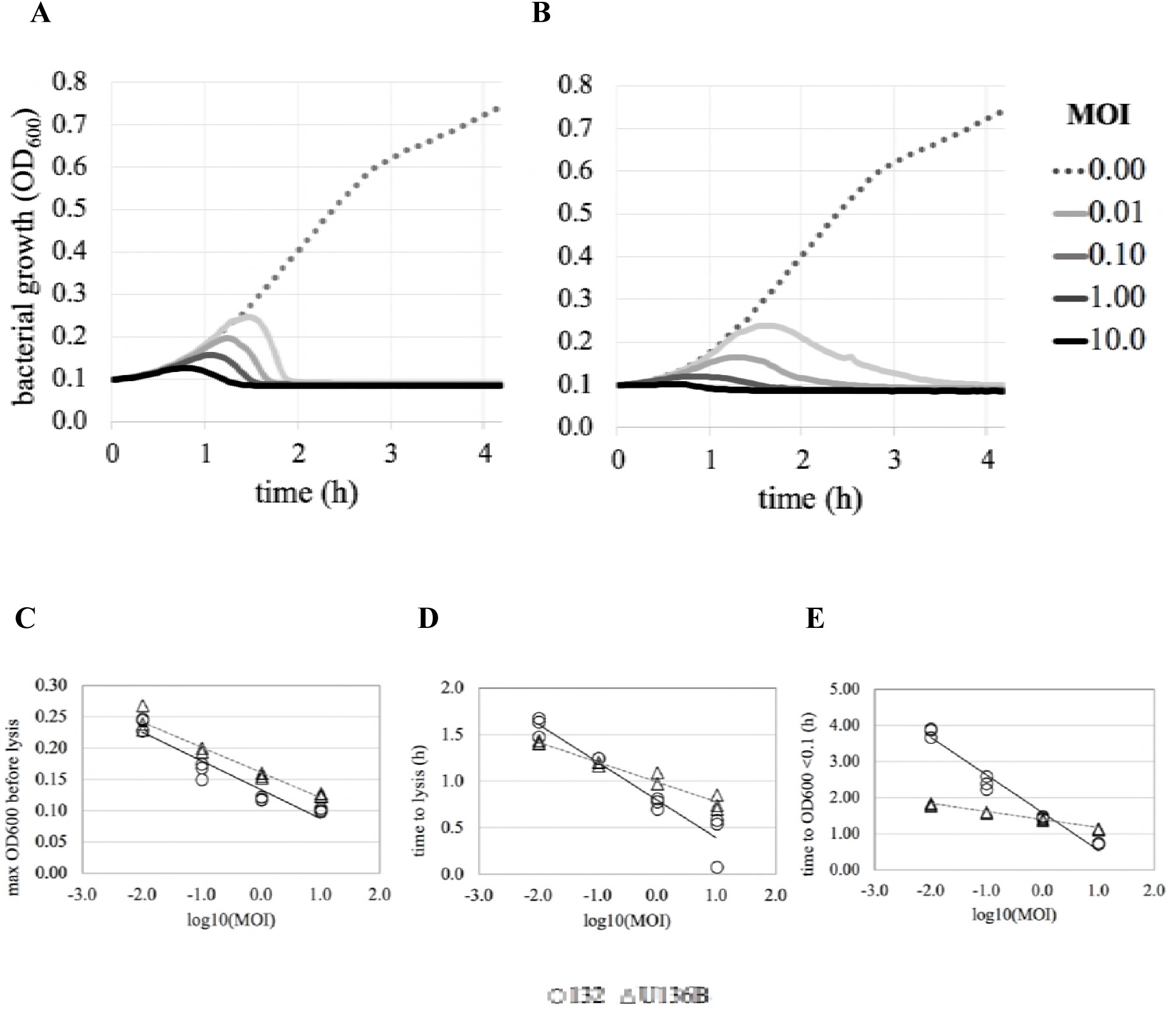
Bacterial lysis at different multiplicities of infection (MOI, the ratio of phage to bacteria) of the *tolC*-dependent phages U136B (Panel A) and 132 (Panel B). Phage addition to bacterial cultures results in cell lysis, with increasing rates of lysis at higher multiplicities of infection (MOI). Each curve shows the mean of three replicates. Panel C) The maximum OD reached before the onset of bacterial lysis, Panel D) The time on lysis onset, and Panel E) The time to complete lysis. Increasing MOI significantly decreased the maximum OD, the time of lysis onset, and the time to complete lysis (p<0.00001 all cases, Table S2). Phage type significantly affected the maximum OD and the complete time to lysis (p<0.01 both cases) but not the time of lysis onset (p = 0.15) (Table S2).

### Phage-resistant mutants

Our EOP and bacterial lysis data suggested that phage resistance in *tolC*^+^ bacterial populations might come about by deletion or modification of the bacterial *tolC* gene. To test whether bacteria can obtain phage resistance mutations in *tolC*, we conducted a selection experiment using phage and bacteria on the surface of agar plates (7). Ten independent bacterial cultures (each seeded from a unique colony of the wild-type bacteria, BW25113) were exposed on plates to phage U136B or 132, and surviving bacterial colonies were readily obtained (Table 2). We randomly picked one colony from each independent culture for further characterization. We also picked additional, non-random colonies that appeared to have unique morphologies, such as the mucoidy phenotype (commonly caused by excess exopolysaccharide production that limits phage infection) (21). We isolated each mutant using the double-isolation technique, then checked for resistance using the cross-streak method. In some cases, the isolated cultures were not resistant to the phage that they had been selected on (Table 3). In those cases, we non-randomly chose another colony from the same independent bacterial culture to complete a set of 10 independent, phage-resistant mutants selected on each phage. We used those sets of 10 independent mutants for further characterization including cross-resistance estimates and sequencing.

**Table 2.** Selection of phage-resistant mutants. Independent cultures of *E. coli* BW25113 were plated at MOI of 1 (phage U136B) or 10 (phage 132).

| Selecting Phage | Surviving colony frequency | Cross resistance to the other phage<sup>1</sup> |
| --- | --- | --- |
| Phage 132 | 2.75 x 10<sup>-7</sup> | 8/10 |
| Phage U136B | 3.20 x 10<sup>-6</sup> | 10/10 |

**Table 3.** Phage-resistant *E. coli* mutants. ‘Culture’ indicates the independent culture the selection was performed on. ‘Selected on’ indicates the phage used for selection. ‘Type’ indicates whether the culture was chosen randomly or nonrandomly from the selection plate.

| Strain | Culture | Selected on | Colony Morphology | Type | Phage 132<sup>R</sup> | U136B<sup>R</sup> | *tolC* Sequenced |
| --- | --- | --- | --- | --- | --- | --- | --- |
| RGB-034 | 1 | U136B | Mucoid | Nonrandom | R | R |  |
| RGB-036 | 1 | U136B | S translucent | Random | R | R | * |
| RGB-040 | 2 | U136B | L opaque | Random | IS | R | * |
| RGB-045 | 3 | U136B | L opaque | Random | IS | R | * |
| RGB-049 | 4 | U136B | S opaque | Random | R | R | * |
| RGB-050 | 4 | U136B | Mucoid | Nonrandom | R | R |  |
| RGB-058 | 5 | U136B | L opaque | Random | IS | R | * |
| RGB-060 | 6 | U136B | S opaque | Random | IS | R | * |
| RGB-065 | 7 | U136B | S opaque | Random | R | R | * |
| RGB-071 | 8 | U136B | S translucent | Random | IS | R | * |
| RGB-074 | 9 | U136B | S opaque | Random | R | IR | * |
| RGB-079 | 10 | U136B | S opaque | Random | IS | R | * |
| RGB-082 | 1 | 132 | L | Random | R | R | * |
| RGB-083 | 1 | 132 | S | Nonrandom | S | S |  |
| RGB-084 | 2 | 132 | L | Random | R | R | * |
| RGB-085 | 2 | 132 | S | Nonrandom | S | S |  |
| RGB-086 | 3 | 132 | S | Random | S | S |  |
| RGB-087 | 3 | 132 | L | Nonrandom | R | R | * |
| RGB-088 | 4 | 132 | L | Random | R | R | * |
| RGB-089 | 4 | 132 | S | Nonrandom | S | S |  |
| RGB-090 | 5 | 132 | L | Random | R | S | * |
| RGB-091 | 5 | 132 | S | Nonrandom | IR | S |  |
| RGB-092 | 6 | 132 | L | Random | IR | R | * |
| RGB-093 | 6 | 132 | S | Nonrandom | R | S |  |
| RGB-094 | 7 | 132 | S | Random | S | S |  |
| RGB-095 | 7 | 132 | L | Nonrandom | R | R | * |
| RGB-096 | 8 | 132 | S | Random | S | S |  |
| RGB-097 | 8 | 132 | L | Nonrandom | R | R | * |
| RGB-098 | 9 | 132 | L | Random | R | R | * |
| RGB-099 | 9 | 132 | S | Nonrandom | IS | R |  |
| RGB-100 | 10 | 132 | S | Random | S | S |  |
| RGB-101 | 10 | 132 | L | Nonrandom | R | R | * |

S: small colony size
L: large colony size S: susceptible (complete clearing of bacterial growth along line of phage, few resistant colonies may be visible) R: resistant (line of bacterial growth looks same before/after/on line of phage) IS: intermediate susceptible (growth is much less dense but still visible after line of phage, more than a few resistant colonies) IR: intermediate resistant (growth is a bit less dense after line of phage but does not look as confluent as totally resistant)

### Cross-resistance to phages

We found that selection by phage U136B yielded cross-resistance to phage 132 (10/10 cases, Table 3). Conversely, selection by phage 132 yielded partial cross-resistance to phage U136B (8/10 cases, Table 3).

### Mutant *tolC* sequences

We found that three of our ten U136B-resistant mutants had mutations in *tolC* (Table 4). None of the phage 132-selected mutants had mutations in *tolC*.

**Table 4.** List of *tolC* mutations identified by targeted Sanger sequencing in mutants resistant to phage U136B. No *tolC* mutations were identified in mutants resistant to phage 132. Location indicates the mutation position from the start of the *tolC* gene.

| U136B<sup>R</sup> Mutant | Mutation Type | Description |
| --- | --- | --- |
| RGB-040 | C472T | premature stop codon (coding 157/493 amino acids) |
| RGB-045 | ΔA at position 1092 | frame shift |
| RGB-071 | ~1.3 kb insertion | IS5 insertion |

### Analysis of phage growth on *omp* gene knockouts

Given that phage 132 could grow on the *tolC* knockout in solid media (albeit, with low efficiency) and lyse bacteria in liquid culture (Figs. 2-3), we hypothesized that it might indirectly interact with *tolC*. Alternatively, it might directly interact with TolC as well as another OMP as an alternative receptor. To test this, we determined efficiency of plating for phage 132 on a variety of OMP gene knockouts. We specifically chose to screen nine knockouts of OMPs known to serve as other phage receptors (14). To increase the resolution of this assay, we used top layer with 50% the normal amount of agar, which we find yields more and larger plaques of phage 132. We found an overall effect of gene knockout on the EOP for phage 132 (F_9,20_ = 58.0, p < 0.0001, Table S3, Fig. 4B). The test also qualitatively replicated the initial result that phage 132 has reduced EOP on the *tolC* knockout, noting that EOP is greater in this modified agar environment (see Methods). Strikingly, the EOP was even further reduced by a single gene knockout of *ompF*, and to a lesser extent, the *ompF* positive regulatory gene *ompR* (Fig. 4B), suggesting that phage 132’s receptor is OmpF.

**Figure 4.**
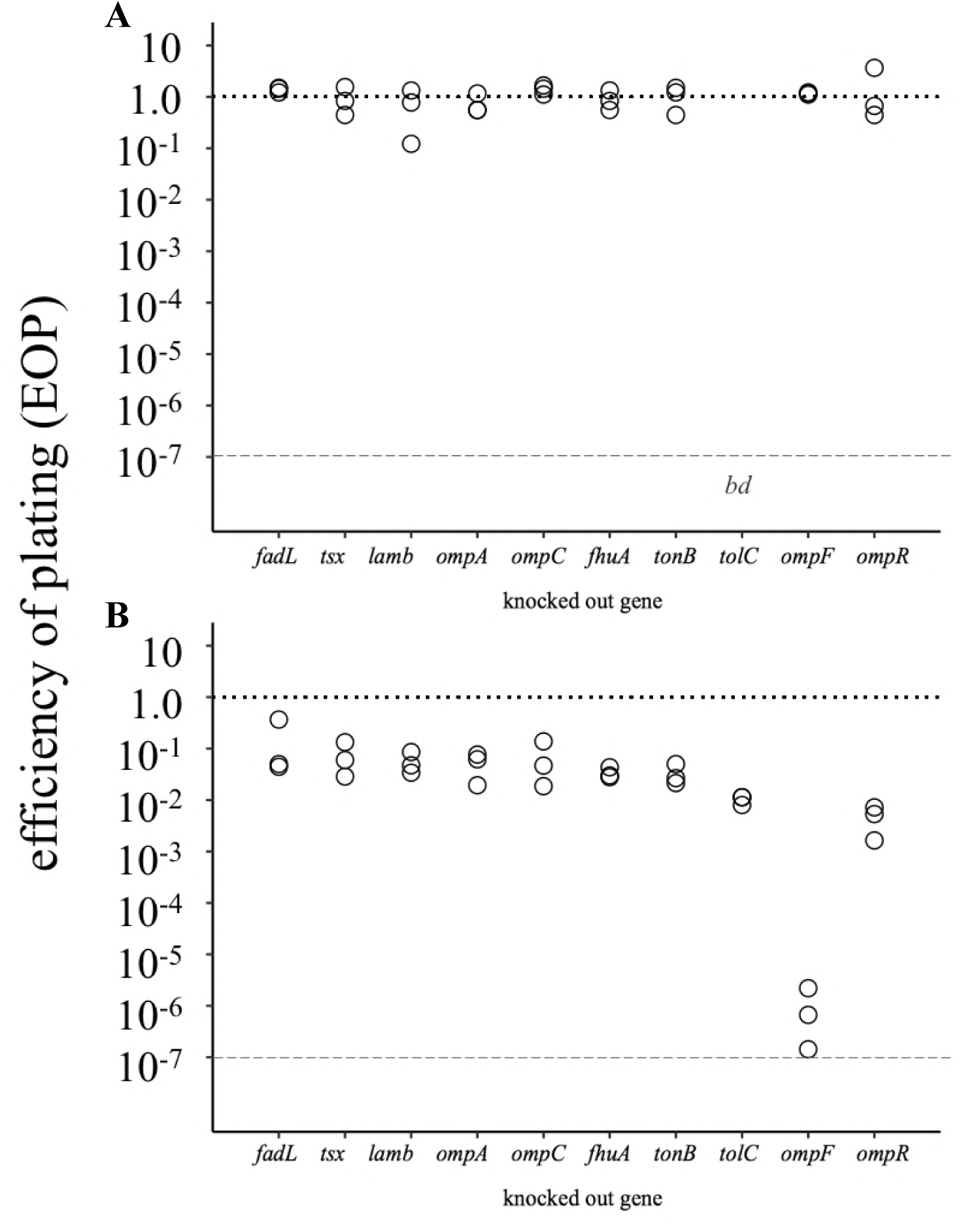
Efficiency of plating of phages U136B (Panel A) and 132 (Panel B) on outer membrane protein-coding gene knockouts. A phage that grows the same (produces an equal number of plaques) on a knockout as on wild-type bacteria has EOP of 1.0 (dotted line). Each assay was conducted in triplicate biological replicates (each replicate phage stock grown independently from an isolated plaque of 132), with technical replicates performed in duplicate. Each point shows the mean of the two technical replicates. A “bd” below the lower, dashed line indicates that the efficiency of plating was <u>b</u>elow the limit of <u>d</u>etection (∼10^-7^). The gene knockout significantly affected the EOP for phage 132 (F9,20 = 58.0, p<0.00001) but not for phage U136B (F8,18, p=0.68) (Table S3).

Observing that phage 132 was affected by more than one OMP gene knockout, we then tested U136’s plaquing efficiency on the same set of genes. U136B was unaffected by knocking out any of the outer membrane protein genes except for *tolC* (Fig. 4A).

### Phage dependence on lipopolysaccharide synthesis genes

To test whether phage U136B and 132 may also rely on lipopolysaccharide (LPS) as primary receptors, we screened for efficiency of plating on knockouts of *rfa* genes, which are involved in LPS synthesis and modification. As expected, we found that knocking out single *rfa* genes greatly impacted efficiency of plating (Fig. 5). Neither phage formed any plaques on four of the knockouts (*rfaC, rfaD, rfaE, rfaP*). Phage 132 uniquely did not form plaques on three additional knockouts (*rfaF, rfaG*, and *rfgH*). For the eight remaining *rfa* knockouts for which EOP could be quantified for both phages (*rfaB, rfaI, rfaJ, rfaL, rfaQ, rfaS, rfaY, rfaZ*), the efficiency of plaquing was significantly effected by both the phage type (F_1,44_ = 21.1, p<0.0001) and *rfa* knockout (F_8, 44_ = 4.7, p<0.001) (Fig. 5, Table S4). Together, these results suggest that phages U136B and 132 use LPS receptors, and that the two phages have different LPS specificity.

**Figure 5.**
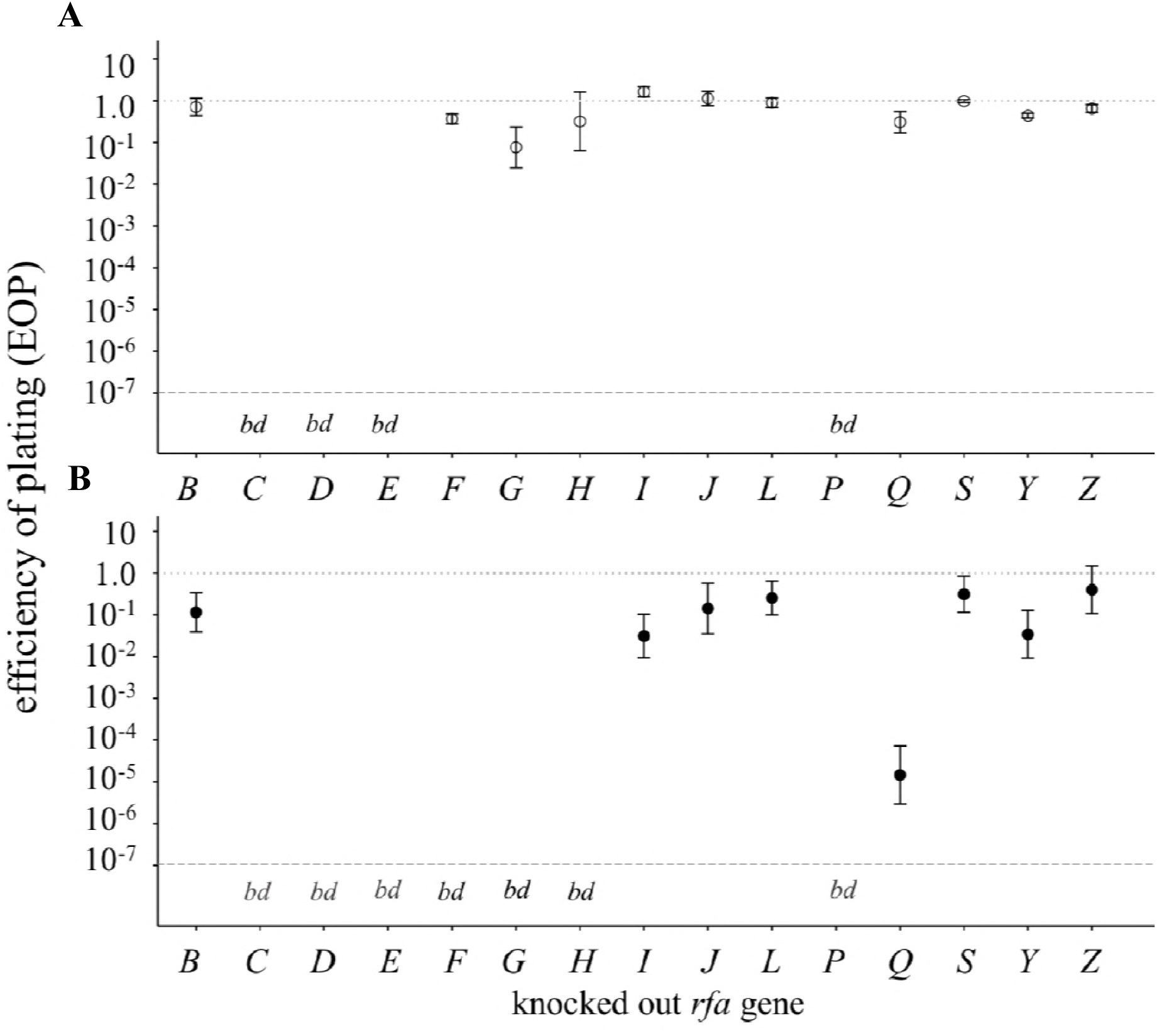
Role of lipopolysaccharide synthesis genes on replication of phages U136B (upper panel) and 132 (lower panel). A phage that grows the same (produces an equal number of plaques) on a knockout as on wild-type bacteria has EOP of 1.0 (upper, dotted line in each panel). Each assay was conducted in triplicate biological replicates (each replicate phage stock grown independently from an isolated plaque). A “bd” below the lower, dashed line within each panel indicates that the efficiency of plating was <u>b</u>elow the limit of <u>d</u>etection (∼10^-7^). For phage-bacteria combinations that were above the detection limit, efficiency of plating was influenced by both the phage (F_1,39_ = 22.4, p<0.0001) and *rfa* gene knockout (F_7,39_ = 4.5, p<0.001) (Table S4).

## Discussion

The interactions between bacteriophages, antibiotic-resistant bacteria, and antibiotic-sensitive bacteria remain unclear. In part, this lack of ecological and evolutionary knowledge is because few phages have been tested for interactions with antibiotic resistance genes. To gain insight into such relationships, we screened a collection of 33 environmental and commercial *E. coli* phages for their inability to infect cells that lacked the antibiotic efflux pump gene *tolC*. The screen identified two environmental phages, U136B and 132, which were recently isolated from a Connecticut swine farm (Table 1). In spatially-structured solid media on *tolC*^+^ bacteria, phage U136B forms easily-visible, medium-sized plaques while phage 132 forms smaller, more difficult-to-enumerate plaques. When *tolC* is absent, phage U136B completely fails to form plaques, while phage 132 forms plaques at reduced efficiency (see Results, ‘*Phage interactions with* tolC^−^ *bacteria’*). The two phages are also morphologically distinct from one another, belonging to different phage families (Fig. 1).

Other than the previously-described phage TLS, to the best of our knowledge these are the only two other *E. coli* phages shown to rely fully or partially on the multidrug efflux gene *tolC* (11, 14). U136B has similar morphology to TLS, a curly-tailed *Siphoviridae* (11). Among other bacterial species, we know of only one phage, OMKO1, that relies on a *tolC* homolog, infecting *oprM*^+^ strains of *Pseudomonas aeruginosa* (5). Despite this apparent rarity of *tolC*-dependent phages, we readily found two out of a modest-sized phage collection of 33 phages. We expect that other environmental phages also rely on *tolC*, and that their absence from the literature reflects under-characterization, rather than low natural abundances. In particular, we expect that future screens of phages from sources where antibiotics are used will yield especially high rates of *tolC*-dependent phages.

While both U136B and 132 both had at least partial reliance on *tolC*, they differently affect lysis and selection in bacterial populations. In liquid culture, U136B rapidly lyses *tolC*^+^ bacteria and has no effect on *tolC*^−^ bacteria (Fig. 2A). Phage 132 efficiently lyses both *tolC*^+^ and *tolC*^−^ bacteria in liquid culture (Fig. 2B). Increasing the multiplicity of infection increases the rate of bacterial lysis for both phages, although the rate of lysis is greater for U136B than phage 132 (Fig. 3). These patterns suggest that phages U136B and 132 might affect bacterial communities in different ways, either by changing their ecological structure (e.g., population sizes, growth rates, death rates, etc.) or by impacting selection on bacteria (e.g., differential survival of phage-sensitive and phage-resistant cells).

To test how each phage might impact selection of phage-resistance mutations, we characterized mutants resistant to each phage. We expected that the phages would select for mutations in the *tolC* gene, as observed previously for *tolC*-dependent phage TLS (11). We observed *tolC* mutations in 3/10 U136B-selected mutants (Table 4). This result is consistent with phage TLS selection experiments, which found that half of TLS^R^ mutants contained *tolC* mutations (11). No *tolC* mutations were observed in phage 132-selected mutants, although we did observe cross resistance between the two phages (Table 3).

Together, the lysis and selection experiments suggest that U136B and 132 interact with the *tolC* gene *via* different mechanisms. Phage U136B’s completely restricted growth on *tolC*^−^ bacteria and *tolC* resistance mutations suggest that it uses TolC as its outer membrane protein receptor. However, we have not yet verified this idea using adsorption assays, complementation tests, or other direct methods. In contrast to U136B, phage 132 seems to interact with *tolC* indirectly. While looking for additional genes required for phage 132 replication, we found that it has severely reduced EOP on *ompF*^−^ bacteria, and to a lesser extent, *tolC*^*−*^ and *ompR*^−^ bacteria (Fig. 4). As both TolC and OmpR increase *ompF* expression (19, 22, 23), one possibility is that phage 132 uses OmpF as an outer membrane receptor. However, at this time it remains unclear whether phage 132 directly interacts with the TolC protein, or indirectly interacts with TolC *via* OmpF or some other mechanism. It is also possible that phage 132 can use either TolC or OmpF as outer membrane protein receptors.

Two other *E. coli* phages (TuIa and T2) use OmpF as a receptor, and several phages can use multiple receptors. Phage T2 can use either OmpF or FadL as a receptor (14). To compare phage 132 to a known multi-receptor, OmpF phage, we tested the EOP of phage T2 on knockouts of *fadL, ompF*, and *ompR* and found only slight reductions in its EOP on either of its receptor knockouts (Fig. S4), while the *ompR* knockout had the greatest reduction in EOP. We hypothesize that this is because ompR affects the regulation of both *fadL* as well as *ompF* (24), thereby reducing the total number of T2 receptors on the cell surface. In contrast, phage 132 appears to be more dependent on *ompF* (Fig. 4B), and thereby more dependent on *tolC*. In addtion to protein receptors, phages U136B and 132 may also require a primary receptor, such as lipopolysaccharaide (LPS) (11, 25, 26). Given that U136B and 132 appear to share some properties with lipopolysaccharide (LPS)-dependent phages TLS and T2, we reasoned that might may rely on LPS synthesis and modifcation genes, such as those in the *rfa* locus. While many other genes are also involved in LPS production, previous work with phage TLS found that half of phage-resistant mutants had mutations in the *rfa* locus (11), and so we used these genes here. Our screen on 13 *rfa* knockouts revealed that removal of many of these genes severely reduced plating efficiency for phage U136B, 132, or both (Fig. 4), with the two phages responding differently to the deletion of various genes. These results suggest that both U136B and 132 have LPS-dependent infection, and that these phages may have unique O-antigen requirements. More importantly, LPS-dependence may provide an easy way for bacteria to evolve resistance to phages U136B and 132, as mutations to *rfa* (or other LPS-related) genes could reduce or eliminate phage infection. This might explain why few of our phage-resistant mutants contained *tolC* mutations (Table 4).

Together, our results show that the antibiotic resistance gene *tolC* can modify bacteria-phage interactions in multiple ways, potentially through a direct interaction as the phage receptor (as we hypothesize for phage U136B) or by modifying expression of the outer membrane protein phage receptor (as we hypothesize for phage 132).

### Conclusion and Future Directions

Our goal here was to identify bacteriophages that rely on the *E. coli* outer membrane protein TolC, which confers multi-drug resistance as part of the AcrAB-TolC efflux pump. Our two newly-isolated phages, U136B and 132, have unique morphologies and differentially impact bacterial populations, including lysis dynamics and selection for phage resistance. Together, these phages will be useful for future studies of evolutionary trade-offs between phage resistance and antibiotic resistance. These phages will also be useful for studying the specific biochemical interactions between phages and their hosts, including whether phage U136B uses TolC and phage 132 uses OmpF for outer membrane attachment. A potential further possibility is the development of these phages for practical application, where they might help to restore drug sensitivity.

## Methods

### Bacterial growth conditions and media

We grew bacteria in LB broth with 10 g tryptone, 5 g yeast extract, and 10 g/L NaCl. LB agar included 15 g/L agar and LB top agar included 7.5 g/L agar unless otherwise noted. Overnight culture incubation was performed at 200 RPM shaking at 37°C.

### Phage library screen

We screened 33 phage isolates collected previously from various sources. We screened each isolate using the plaque spot test on host lawns of wild-type and knockout *E. coli* from the Keio collection obtained from the Yale Coli Genetic Stock Center (Table 1).

### Efficiency of plating

Those phages for which a difference in plaquing was observed in the library screen were then screened for differences in efficiency of plating (EOP, the number of plaques formed on a mutant relative to wild-type bacteria). We determined the initial EOP for phages U136B and 132 on the *tolC* knockout using a full plate dilution series in standard top agar (7.5 g/L agar). For all other EOP assays, we used we used top agar that contained 3.8 g/L, which is 50% the typical amount of agar, and serial dilution spot tests of 2 µl to obtain phage titers on each bacterium. (We found that phage 132 generally formed small, difficult-to-count plaques, were more easily enumerated in the 3.8 g/L formula.)

### Chloroform/vortex sensitivity assay

We tested phage sensitivity to vortexing and to chloroform exposure under the typical conditions we use during phage preparation. We subjected test phages to three treatments: LB only, LB plus vortexing at moderate speed (setting 4/10 on a Scientific Industries Vortex Genie 2) for 3 seconds, and LB with 1% v/v chloroform. We plated a dilution series of 1.5-µL spots of phage on LB agar before and after each treatment in triplicate.

### Mutant selection procedure

To select for phage-resistant bacteria, we mixed wild-type bacteria and phages, then spread plated them onto LB agar and incubated overnight at 37°C. To ensure the isolation of independent mutations, each replicate was grown from a single isolated colony. To confirm the multiplicity of infection (MOI, the ratio of phage particles to bacterial cells) on the agar plate, we plated bacteria and phage in triplicate and counted CFU/ml and PFU/ml, respectively. From each mutant-selection plate, we picked a random colony, plus any colonies of notable morphology, and streaked each onto LB agar plate and incubated at 37°C overnight. We re-streaked the colonies to obtain isolates and grew each in 10 mL LB at 37°C with shaking overnight. We archived freezer stocks of each mutant in 20% glycerol, stored at −80°C.

### Mutants’ phage resistance and cross-resistance

We checked for resistance and cross-resistance using the cross-streak method (27), modified as follows. A line of 10 µl of high-titer phage stock was placed along an LB agar plate. When the phage streak had dried, 1 µl of bacterial culture was streaked across the phage. Cultures with no growth past the phage line were scored as phage-susceptible. Cultures with equal growth past the phage line were scored as phage-resistant. Cultures with decreased growth past the phage line were scored as intermediate-resistant.

### Bacterial growth curves

Cultures of wild type and Δ*tolC* were grown to exponential phase as measured by optical density at 600nm wavelength (OD_600_) of 0.25-0.38. The exponential-phase cultures were diluted 1:5 into wells with LB to a total volume of 200 µl with phage to reach the target multiplicities of infection (MOI, the ratio of phage to cells). Cultures were incubated at 37°C with shaking at 288 RPM for 18 h and OD_600_ read every 2 minutes using an automated spectrophotometer (TECAN microplate reader).

### TEM

Transmission electron micrographs were collected on high-titer phage lysates at the Yale Electron Microscopy facility in the Center for Cellular and Molecular Imaging. Samples were negatively stained with 2% uranyl acetate and imaged with FEG 200kV transmission EM.

### PCR and Sanger sequencing

We extracted DNA from overnight cultures grown in LB using the Qiagen DNeasy Blood and Tissue Kit (Cat. # 69506), following the product’s protocol for Gram-negative bacteria and the optional RNAse step. We amplified the *tolC* gene using Promega GoTaq DNA polymerase (Cat. # M829) with standard buffer and primers AJL-001 and AJL-002 (Table S1), with an annealing temperature of 54°C. Sanger sequences were collected by the DNA Analysis Facility on Science Hill at Yale using all four primers (Table S1). We aligned sequences to the reference sequence BW25113 (Table 1) using SeqMan Pro (LaserGene, DNASTAR, Madison, WI), confirming mutations by assessing the sequence trace data. We determined the identity of the insertion in the case of RGB-071 using BLAST (28) via the NCBI Nucleotide online tool (https://blast.ncbi.nlm.nih.gov/Blast.cgi) against reference sequence CP009273.1 for *E. coli* strain BW25113 (Table 1).

### Statistical analysis

We used linear regression to test the effect of log-transformed MOI and phage type on bacteria lysis time parameters. We used ANOVAs to test the effect of bacterial *omp* gene knockouts on log-transformed efficiency of plating (EOP) data. (We conducted these analyses separately for phage 132 and U136, as the two phages were tested in separate experiments and were thereby potentially subjected to block effects.) To test the effects of bacterial *rfa* gene knockout and phage type on EOP, we used an ANOVA with both gene and phage type independent variables. All analyses were performed in R (29) using custom scripts.

## Acknowledgments

Our work was supported by NSF Cooperative Agreement DBI-0939454 through the BEACON Center for the Study of Evolution in Action, Howard Hughes Medical Institute Campus Grant #52008128, a Yale College Dean’s Office Science Research Fellowship to RGB, and a Yale Trumbull College Richter Fellowship to AF. We thank Caroline Turner, Mike Wiser, and Mike Blazanin for comments on the manuscript, and Mike Blazanin, Jennifer Yuet Ha Lai, Padma Mamillapalli, Geetha Suresh, and Vicki Taccardi for assistance in the lab.

